# Defining Predictors of Successful Early Career to Independent Funding Conversion Among Surgeon-Scientists

**DOI:** 10.1101/2022.03.30.486442

**Authors:** Sonya S. Keswani, Walker D. Short, Steven C. Mehl, Kavya L. Singampalli, Umang M. Parikh, Meghana Potturu, Leighanna Masri, Oluyinka O. Olutoye, Lily S. Cheng, Alice King, Timothy C. Lee

## Abstract

**Introduction:** The National Institutes of Health (NIH) provides research funding to scientists at different stages of their career through a range of grant awards. Early-stage researchers are eligible for mentored Career Development (K) awards, to aid in the transition to independent NIH funding. Factors such as education, subspecialty, and time to funding have been studied as predictors of obtaining independent awards in nonsurgical specialties. However, in surgery, the importance of these factors has yet to be clearly elucidated. We aim to identify predictors of K to independent award conversion among surgeon-scientists to understand how to better support early-stage researchers transitioning to independent careers.

**Materials and Methods:** In July 2020, the NIH Research Portfolio Online Reporting Tools database was queried for individuals affiliated with surgery departments who received NIH Career Development Awards (between 2000 and 2020). The following factors were analyzed: publications, institution, degrees, year of completion of training, and gender.

**Results:** Between 2000 and 2020, 228 surgeons received K Awards, of which 44% transitioned to independent funding. On average, surgeons received a K award 4.0 years after completing fellowship training and an independent award 5.4 years after receiving a K grant. The time to receiving a K award was predictive of successfully achieving independent funding, and those with independent funding had a significantly greater number of publications per year of their K-award.

**Conclusion:** Surgeons successful in transitioning to independent NIH awards do so approximately 9 years after finishing fellowship. Publication track record is the main factor associated with successful conversion from a K award. Surgery departments should emphasize manuscript productivity and develop strategies to minimize time to independent funding to help K-awardees begin independent research careers.

## Introduction

Academic surgeon-scientists have a unique role in impacting the advancement of medical knowledge, bringing ideas from the operating room to the bench, and in turn translating that research back to the forefront of surgical care. From Dr. Alexis Carrel’s Nobel Prize-winning work on organ transplantation to Dr. Stanley Dudrick’s development of total parenteral nutrition, surgeons have a history of massively impactful contributions to the science of medicine.^1,2^ Given the reliance of academic scientists on funding from the National Institutes of Health (NIH), complicated by the continued diminution of NIH funding,^3^ it is imperative that we understand the predictors of success of those surgeon-scientists who are successfully able to achieve independent NIH funding. In 1972, the NIH commenced a new category of funding termed Career Development Awards (CDAs), or K-series awards. The stated objective of K-awards is “to help ensure that a diverse pool of highly trained scientists are available in adequate numbers and in appropriate research areas to address the Nation’s biomedical, behavioral, and clinical research needs.”^4^ Since its inception, the K-to-R-award transition has been a commonly pursued means to research funding independence, however surgeons tend to be much less likely to pursue NIH CDA’s compared to their non-surgical colleagues.^5^

Surgeon-scientists have historically been underrepresented as NIH grant awardees in the United States, receiving both fewer and less valuable rewards compared to non-surgical physician scientists.^6^ A recent uplifting update by Demblowski and colleagues demonstrated an increase in the proportion of funded surgeons over the past decade, with maintenance of R-series funding in that time period.^7^ Despite this encouraging improvement in surgical funding and the far-reaching impact that surgeon-scientists have made in advancing medical therapies, the proportion of surgeons receiving funding through the NIH does not reflect the number of clinically active surgeons in the workforce.^8^ This inconsistency can be explained by the known data that surgeon-scientists apply for fewer grants compared to nonsurgical investigators and is exacerbated by the lack of surgeon representation on NIH study sections.^9,10^ Interestingly, the publication “return on investment” of National Institute of General Medical Sciences (NIGMS) funding is almost 5-fold greater for a first time NIH grant awardee compared to a senior investigator receiving a third R01,^11^ which even further highlights the importance of funding enthusiastic early-career surgeon-scientists. Although many remedies to improve the funding of surgeons has been proposed, limited analysis of the factors which predict successful transition to independent funding has been performed.

Herein, we use publicly available data to examine the factors such as gender, degree status, type of research, and number of publications which are associated with successful transition from mentored to independent funding. We hypothesize that the number of peer-reviewed publications associated with surgeon’s K-funding is the principle modifiable factor that predicts successful transition to independent funding.

## Material and Methods

### Study Population

In this study, the population included clinicians with an MD or equivalent degree who completed a surgical residency and received an NIH grant between 2000 and 2020. Institutional and departmental websites were searched to determine clinical training. Funding information for each clinician was determined through the NIH Research Portfolio Online Reporting Tools (RePORT) database. This resource provides information on NIH funding for investigators and reports on projects and research activities. No individuals were directly contacted. To ensure information for each physician was accurate, all recorded results were cross-verified using the institutional affiliation and department listed on NIH RePORT.

### Data Collection and Variables

Publicly available data in the NIH RePORT database was queried in July 2020 for those affiliated with surgery departments and had received NIH Career Development Awards (K01, K08, or K23) between 2000 and 2020. For each surgeon-scientist that met the criteria and received a K01, K08, or K23 award, the dates and type of awards received, publication success (defined as the number of publications listed on NIH reporter under the grant divided by the number of years the grant was active), number and years of independent awards, and total independent NIH funding were recorded. Information pertaining to demographics, including the year of clinical training completion, gender, subspeciality, degrees earned, type of research (basic science versus clinical), and time from completion of training were collected through faculty biography pages. Institutional characteristics were collected and included NIH funding ranking data collected by the Blue Ridge Institute for Medical Research,^12^ type of institution (public versus private), and location (South, West, Midwest, Northeast).

### Statistical Analysis

The primary outcome of interest was successful K-to-R transition. Potential predictors or exposures evaluated included applicant characteristics (gender, additional degrees, surgical specialty, time from completion of training, and type of research), productivity (number of publications/year), institutional characteristics (ranking, public versus private, region), and amount of funding per year. Continuous outcomes were reported as median with interquartile ranges (IQR). Categorical outcomes were reported as percentages. Univariate analysis was performed using two sample Student’s t-test to compare continuous variables and Chi–Square or Fisher’s exact test to compare categorical variables. A multivariable logistic regression model was used to evaluate the association between different predictors and successful K to R transition. Model covariates were selected a priori and included gender, attending time, type of research, type of institution (private or public), number of publications per year, national institutional ranking, and amount of funding per year. Adjusted odds ratio (aOR) and 95% confidence interval (CI) were calculated. A *p* value of <0.05 was considered significant. All analyses were performed using Stata version 16.0 (StataCorp, College Station, TX).

## Results

### Demographics

At the time of the query, we identified 228 surgeons who received K-awards between 2000 and 2020 (**Figure 1**). Overall, 44.7% of these surgeons successfully converted these CDA’s to independent NIH grants (2000 – 2014, 53.6% successful; 2015 – 2020, 20.3% successful). The average time between K-award and R-funding for all successful transitions was 5.05 years. Therefore, to achieve a more accurate analysis of predictors of successful transition, surgeons who received their K-award between 2015 and 2020 (n=60) were excluded from analysis. This left 168 surgeons who obtained a K-award between 2000-2014 for further analysis.

**Figure 1:**
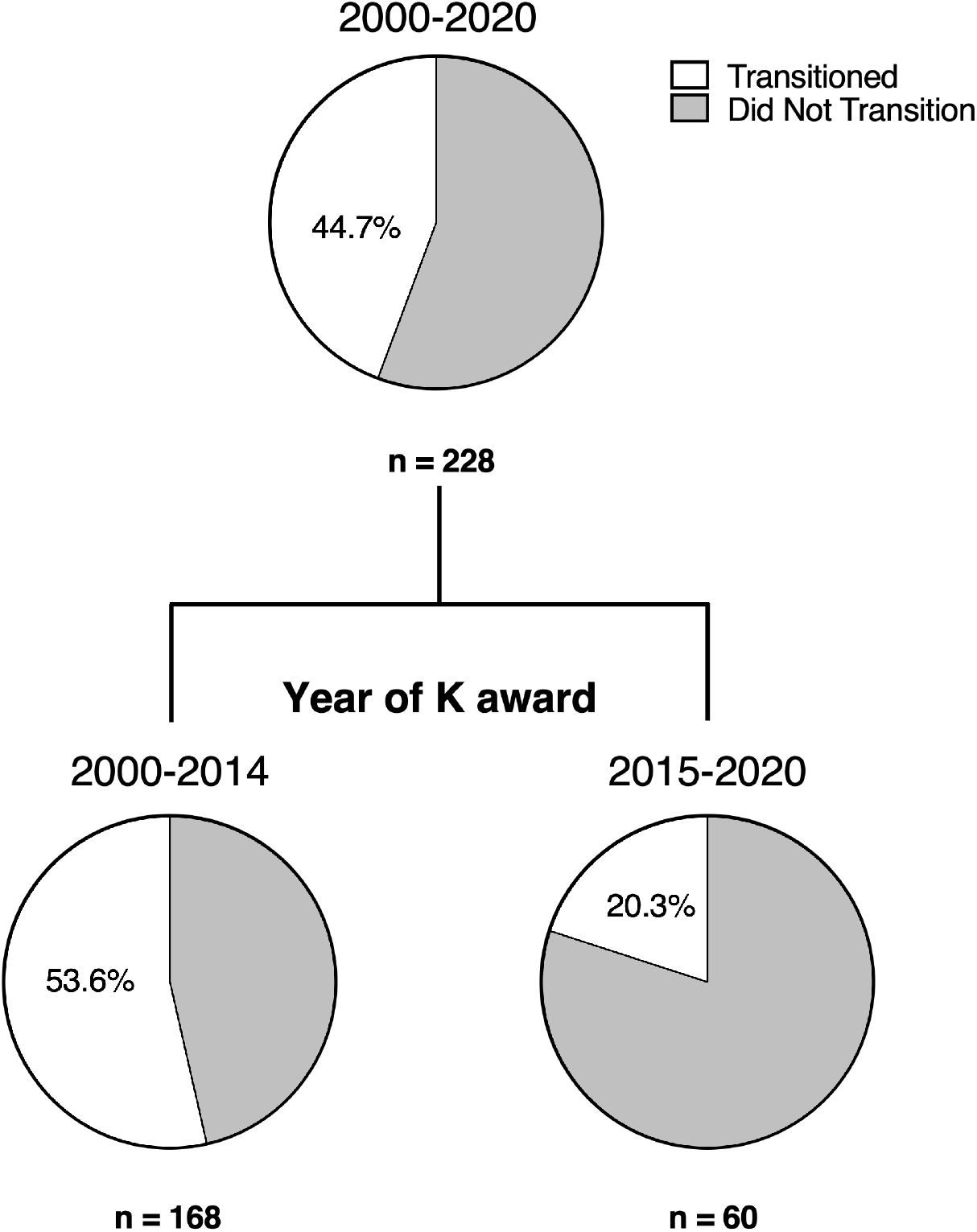
Proportion of K-to-R Transition by Time Period.

Of the remaining K-awardees, 130 were male (77%), 131 (78%) conduct basic science research, and 58 (35%) have additional degrees beyond MD. 16 (9 %) practiced in general or minimally invasive surgery, 29 (17%) in trauma/critical care, 31 (18%) in surgical oncology, 21 (12%) in cardiothoracic surgery, 32 (19%) in vascular surgery, and 19 (7%) in pediatric surgery.

### Predictors of Successful Transition

Applicant characteristics including gender, additional degrees, surgical specialty, and type of research were not significantly associated with successful transitions (**Table 1). Figure 2** displays number of surgeons who had successful and unsuccessful transition when stratified by specialty Less time from completion of training and increased number of publications per year was associated with a successful transition. There was no association between type of research and successful transition to independent funding. Institutional characteristics, including national ranking, public versus private institution, and location were not associated with successful transitions. When controlling for applicant characteristics, productivity, institutional characteristics, and amount of funding per year with multivariate analysis, each paper published per year was associated with a 21% increase (aOR 1.21, 95% CI 1.02 – 1.43) in successful transition, and every additional year after completion of training was associated with a 23% decrease in successful transition (aOR 0.77, 95% CI 0.65 – 0.90) (**Figure 3**).

**Table 1:**
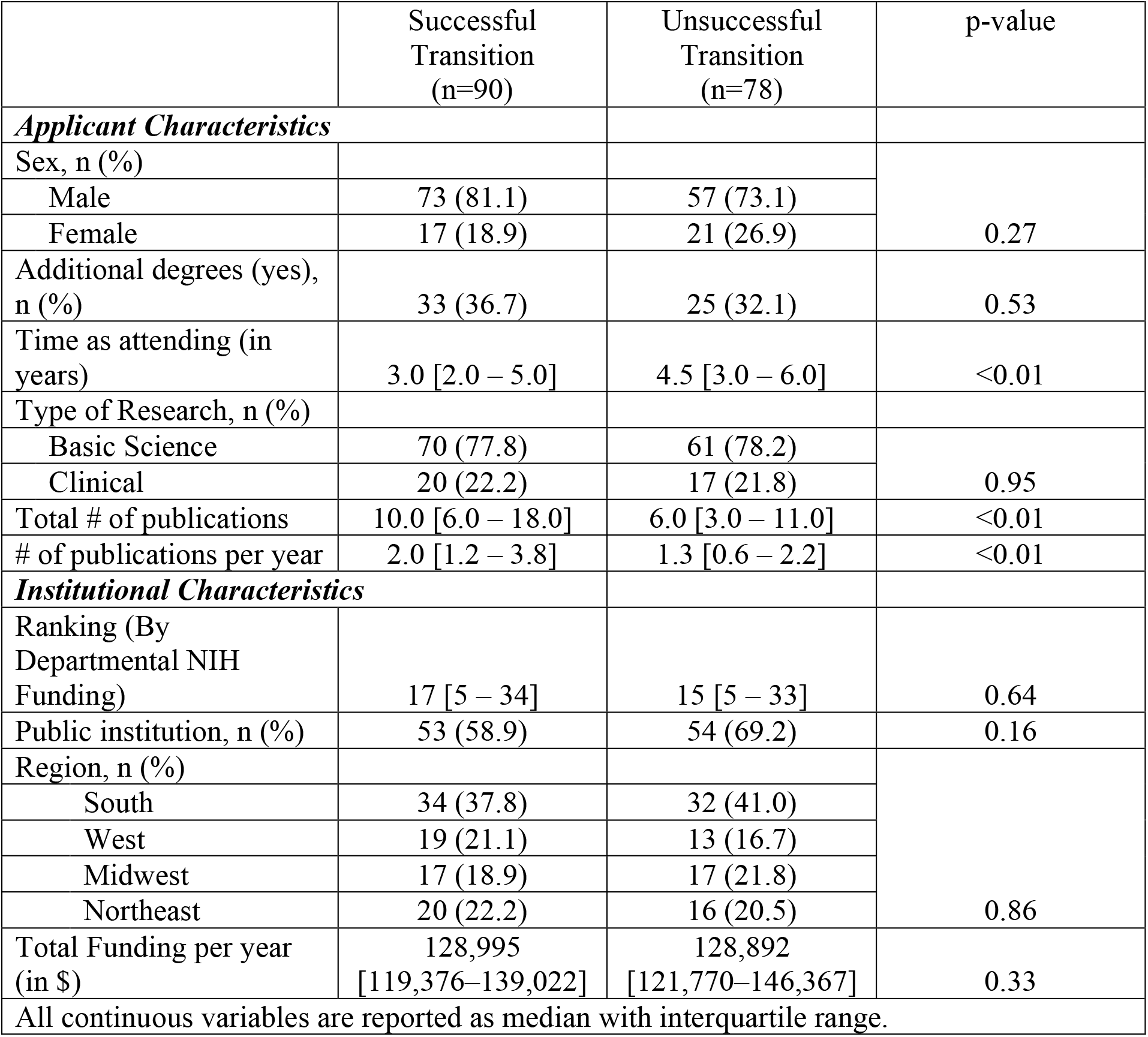
Applicant and Institutional Details stratified by successful transition (2000 – 2014)

**Figure 2:**
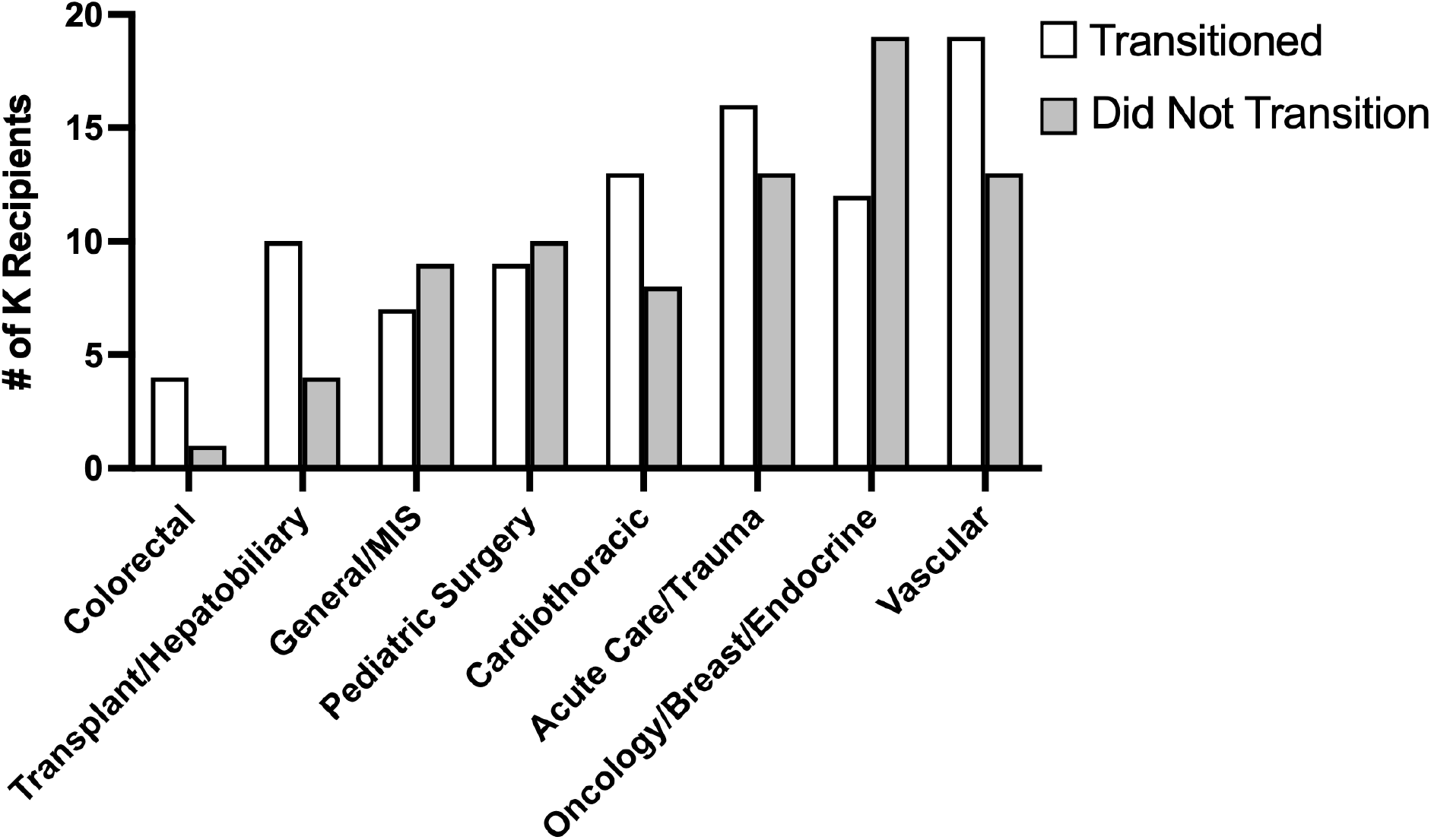
Breakdown of Successful K-to-R Transition Between Surgical Specialties.

**Figure 3:**
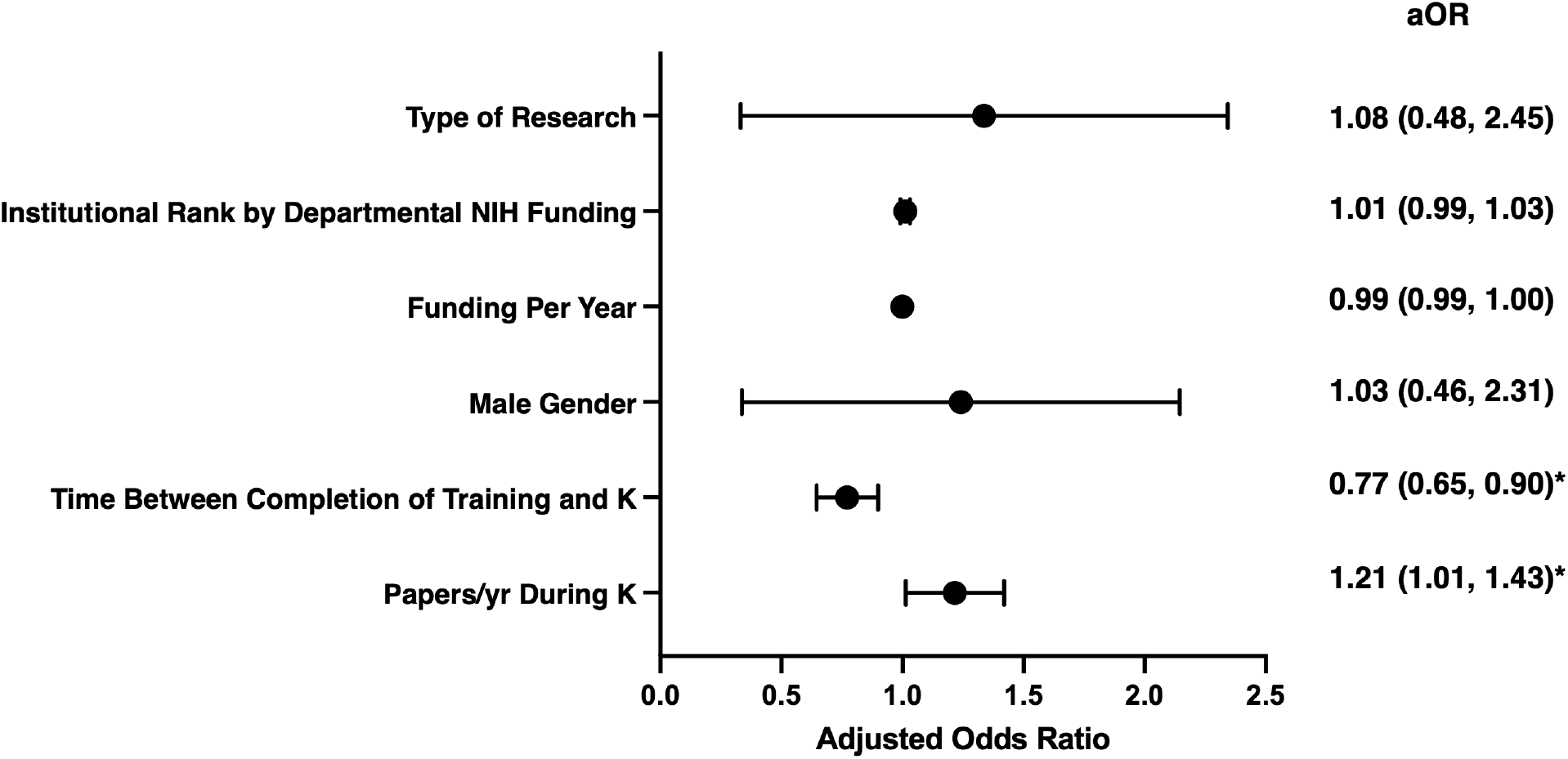
Forest Plot for Predictors of Independent Funding.

## Discussion

Here we have demonstrated that the most critical measurable factor which predicts the successful transition of surgeon-scientists from mentored NIH funding to independence is the quantity of peer reviewed publications resulting from CDA funding. The ability of surgeons to successfully transition to independent funding was irrespective of their subspecialty, type of research, region, gender, and institutional prowess on both univariate and multivariate analysis. Similarly, the amount of funding received from K-awards did not impact a successful transition to independence. Surgeons who successfully obtained mentored awards earlier in their post-training career were more likely to transition to independent funding, indicating the importance of early career protected time and support from institutions.

Given that the most important modifiable factor for early career surgeon-scientists is the quantity of publications resulting from their K-award, the most impactful way home institutions can support these researchers is with generous assistance towards work that may be reportable.??? aimed towards producing publishable work. Balancing early clinical duties with research time, particularly “wet-lab” basic science research, creates an inherent challenge to young surgeons who still do not have the clinical experience and confidence of their more senior partners. This is compounded by the growing concern that United States general surgery residency graduates are becoming less prepared for independent practice following their training.^13^ In a survey of members of the Association for Academic Surgery (AAS) and Society of University Surgeons (SUS), a resounding 68% of respondents felt that it is unrealistic to be successful scientists in the modern hospital environment.^14^ A majority of the basic science researchers polled responded that clinical demands, administrative duties, and desire for work-life balance were prohibitive in their desire to perform research activities. These views are reflected in a significant decrease in NIH career development awards to surgeons over the past 20 years.^15^ Therefore, newly graduated surgeons who crave a significant proportion of their efforts to be devoted to research must have transparent discussions with institutions and departmental heads when considering employment opportunities. With some exceptions, K-awards require that 75% of an awardee’s time be devoted to research activity rather than clinical responsibilities. The salary for this research time is compensated by the K-award, however the maximum salary support from a K-award is $197,300.^16^ Supposing the institution compensates the remaining 25% on the same pay scale this comes to a yearly salary of about $263,000, significantly below the expected starting salary for a general surgeon in most regions of the US. Ideally, the surgeon’s institution is willing to bridge this income gap by more heavily weighting the salary of the clinical time agreed upon. Indeed, even prior to the K-award, some protected research time may be required by the young surgeon to generate preliminary data to be used in the K application, as often the surgeon will not come with unpublished preliminary data from their research time during residency. In all, our data indicates that institutions who prioritize the successful acquisition of NIH funding by their staff surgeons must be willing to do what they can to support their productivity as measured by publications, implying a meaningful effort towards protected research time and salary support. Additionally, clinical demands on young surgeons necessitate institutions and mentors provide other forms of support beyond financial compensation.

With the rapid pace of scientific and methodological advances, it is understandable that many AAS and SUS survey respondents felt that it is unrealistic to expect academic success from surgeons in today’s environment. Basic science research, in particular, requires absolute dedication to become an expert in a particular field. Thus, the idea of appropriate mentorship of a mentor who already has expertise in the area of interest of the young surgeon is essential. While the mentorship aspect of CDA’s is engrained in their mission, a simple letter of support signing off on a K-application, without significant logistical and scientific support, dooms that surgeon to failure in achieving independent funding. Despite the academic prowess tied to private institutions and institutions with the greatest amount of NIH funding for surgical departments, these factors did not predict successful transition to independent funding.

Surprisingly, previous work has shown investigators at the institutions with the least funding tend to be more productive, as measured by publications per NIH dollar.^17^ Given that the sole significant predictor of transition is the number of publications during the K period, this may indicate that the success of the transition to independent funding is determined more by the quality of the mentorship provided and alignment of the scientific focus of the mentor, institution, and awardee. As the title of this grant program implies, exceptional faculty mentorship is a crucial and difficult-to-measure factor for the success of junior academic surgeons.

Women represented around 20% of K-awardees in our cohort, which falls short of the proportion of women in academic surgery overall, roughly 30% as of 2018.^18^ In a longitudinal analysis of clinician-scientists with CDA’s, women were awarded less funding and had fewer publications than men as their careers progressed.^19^ This corresponded to proportionately fewer women who achieved the role of full-professor compared to men. While our data indicates there is no statistical advantage for men to achieve transition to independent funding, the disparity between the genders is clear. With the goal of achieving gender equity in academic surgery, it should be the aim of academic surgeons to encourage more female matriculation into surgical specialties. Female surgeons feel that there is a deficiency of mentorship in their field, and that this poses a hindrance to professional advancement.^20^ Though there is a notable paucity of female surgeons at the professor level, most female surgeons prefer gender-concordant mentorship. While this does not preclude the valuable mentorship which male surgeons can provide, it does add to the importance of advocating for the success and promotion of female surgeons to senior mentorship roles.^20,21^ Beyond continued efforts to eliminate discrimination, mentoring and inspiring more female surgeon-scientists will lead to a feed-forward cycle to increase the pipeline of female medical students involved in surgical research.

The inherent limitation of this study is that only publicly available data was utilized in our analysis. There are many other variables at play, including any internal funding opportunities provided by home institutions, amount of protected research time, and the somewhat unmeasurable quality of the faculty mentor. In particular, grant applicants require considerable administrative support in writing applications for independent funding. Institutions which provide not only assistance with budgeting and formatting, but also insights from senior scientists are likely to have more success in generating successful applications. It must also be noted that some surgeon-scientists may have enough preliminary data to apply directly for R funding, likely removing a proportion of applicants from our data pool who likely would have successfully transitioned from K funding.

## Conclusions

Our data indicate surgeons successful in transitioning to independent NIH awards do so approximately 9 years after finishing fellowship. Publication track record and time from completion of training are the main factors contributing to successful conversion from a K award. Surgery departments should emphasize manuscript productivity and develop strategies to minimize time to independent funding to help K-awardees begin independent research careers.

